# Mini-barcodes are equally useful for species identification and more suitable for large-scale species discovery in Metazoa than full-length barcodes

**DOI:** 10.1101/594952

**Authors:** Darren Yeo, Amrita Srivathsan, Rudolf Meier

## Abstract

New techniques for the species-level sorting of millions of specimens are needed in order to accelerate species discovery, determine how many species live on earth, and develop efficient biomonitoring techniques. These sorting methods should be reliable, scalable and cost-effective, as well as being largely insensitive to low-quality genomic DNA, given that this is usually all that can be obtained from museum specimens. Mini-barcodes seem to satisfy these criteria, but it is unclear how well they perform for species-level sorting when compared to full-length barcodes. This is here tested based on 20 empirical datasets covering ca. 30,000 specimens and 5,500 species, as well as six clade-specific datasets from GenBank covering ca. 98,000 specimens for over 20,000 species. All specimens in these datasets had full-length barcodes and had been sorted to species-level based on morphology. Mini-barcodes of different lengths and positions were obtained *in silico* from full-length barcodes using a sliding window approach (3 windows: 100-bp, 200-bp, 300-bp) and by excising nine mini-barcodes with established primers (length: 94 – 407-bp). We then tested whether barcode length and/or position reduces species-level congruence between morphospecies and molecular Operational Taxonomic Units (mOTUs) that were obtained using three different species delimitation techniques (PTP, ABGD, objective clustering). Surprisingly, we find no significant differences in performance for both species- or specimen-level identification between full-length and mini-barcodes as long as they are of moderate length (>200-bp). Only very short mini-barcodes (<200-bp) perform poorly, especially when they are located near the 5’ end of the Folmer region. The mean congruence between morphospecies and mOTUs is ca. 75% for barcodes >200-bp and the congruent mOTUs contain ca. 75% of all specimens. Most conflict is caused by ca. 10% of the specimens that can be identified and should be targeted for re-examination in order to efficiently resolve conflict. Our study suggests that large-scale species discovery, identification, and metabarcoding can utilize mini-barcodes without any demonstrable loss of information compared to full-length barcodes.

## Introduction

The question of how many species live on earth has intrigued biologists for centuries, but we are nowhere close to having a robust answer. We do know that fewer than 2 million have been described and that there are an estimated 10-100 million multicellular species on the planet (Roskov et al. 2018). We also know that many are currently being extirpated by the “sixth mass extinction” (Ceballos et al. 2015; Sánchez-Bayo and Wyckhuys 2019), with potentially catastrophic consequences for the environment (Cafaro 2015). Monitoring, halting, and perhaps even reversing this process is frustrated by the “taxonomic impediment”. This impediment is particularly severe for “invertebrates” that collectively contribute much of the animal biomass (e.g., Stork et al. 2015; Bar-On et al. 2018). Most biologists thus agree that there is a pressing need for accelerating species discovery and description. This very likely requires the development of new molecular sorting methods because the traditional approach involving parataxonomists or highly-trained taxonomic experts is either too imprecise (Krell 2004) or too slow and expensive. However, any replacement method based on DNA sequences should be accurate, but also (1) rapid, (2) cost-effective and (3) and largely insensitive to DNA quality. These criteria are important because tackling the earth’s biodiversity will likely require the processing of >500 million specimens, even under the very conservative assumption that there are only 10 million species (Stork 2018) and a new species is discovered with every 50 specimens processed. Cost-effectiveness is similarly important because millions of species are found in countries with limited funding and only basic research facilities. On the positive side, many species are already represented as unsorted material in museum holdings, but such specimens often yield degraded DNA (Cooper 1994). Therefore, methods that require DNA of high-quality and quantity are not likely to be suitable for large-scale species discovery in invertebrates.

### High-throughput species discovery with barcodes

Conceptually, species discovery and description can be broken up into three steps. The first is obtaining specimens, the second, species-level sorting, and the third, species identification or description. Fortunately, centuries of collecting have gathered many of the specimens that are needed for large-scale species discovery. Indeed, for many invertebrate groups it is likely that the museum collections contain more specimens of undescribed than described species; i.e., this unsorted collection material represents vast and still underutilized source for species discovery (Lister et al. 2011; Kemp 2015; Yeates et al. 2016). The second step in species discovery/description is species-level sorting, which is in dire need of acceleration. Traditionally, it starts with sorting unsorted material into major taxa (e.g., order-level in insects). This task can be accomplished by parataxonomists but may in the future be guided by robots utilizing neural networks (Valan et al. 2019). In contrast, the subsequent species-level sorting is usually time-limiting because the specimens for many invertebrate taxa have to be dissected and slide-mounted before they can be sorted to species-level by highly-skilled specialists; i.e., the traditional techniques are neither rapid nor cost-effective. This impediment is likely to be responsible for why certain taxa that are known to be abundant and species-rich are particularly poorly studied (Bickel 1999).

An alternative way to sort specimens to species-level would be with DNA sequences. This approach is particularly promising for metazoan species because most multicellular animal species can be distinguished based on cytochrome c oxidase subunit I (*cox1*) barcode sequences (Hebert et al. 2003). However, such sorting requires that every specimen is barcoded. This creates cost- and scalability problems when the barcodes are obtained with Sanger sequencing (see Taylor and Harris 2012). Such sequencing is currently still the standard in many barcoding studies because the animal barcode was defined as a 658-bp long fragment of *cox1* (“Folmer region”: Folmer et al. 1994), although sequences >500-bp with <1% ambiguous bases are also considered BOLD-compliant (Barcode Of Life Data System: BOLDsystems.org). The 658-bp barcode was optimized for ABI capillary sequencers but it has become a burden because it is not suitable for cost-effective sequencing with Illumina platforms.

Due to these constraints, very few studies have utilized DNA barcodes to sort entire samples into putative species (but see Fagan-Jeffries et al. 2018). Instead, most studies use a mixed approach where species-level sorting is carried out based on morphology before a select few specimens per morphospecies are barcoded (e.g. Riedel et al. 2010). Unfortunately, this two-step process requires considerable amounts of skilled manpower and time and does not allow for an unbiased assessment of congruence-levels between morphology and DNA barcodes.

### Obtaining Barcodes with High Throughput Sequencing

Fortunately, scalability and cost-effectiveness are hallmark features of the new high throughput sequencing (HTS) technologies. These technologies are particularly suitable for sequencing the kind of degraded DNA that is typical of museum specimens. Indeed, hybrid capture has already been optimized for the use with old museum specismens (Bi et al. 2013; Guschanski et al. 2013; Blaimer et al. 2016) and is likely to play a major role for the integration of rare species into taxonomic and systematic projects. It will be difficult and expensive, however, to apply enrichment methods to millions of specimens because such methods require time-consuming and expensive molecular protocols (e.g., specimen-specific libraries). Fortunately, for most species the initial species-level pre-sorting can be achieved using barcodes that can be obtained via “tagged amplicon sequencing” on a variety of next-generation sequencing platforms (“NGS barcodes”: Wang et al. 2018; Yeo et al. 2018).

Until recently, obtaining full-length NGS barcodes via tagged amplicon sequencing was difficult because the reads of most next-generation-sequencing platforms were too short for sequencing the full-length barcode. This has now changed with the arrival of third-generation platforms (ONT: MinION: Srivathsan et al. 2018, 2019; PacBio: Sequel: Hebert et al. 2018). These platforms, however, come with drawbacks; *viz* elevated sequencing error rates and higher cost. These problems are likely to be overcome in the future (e.g. Yang et al. 2018), but this would still not solve the main challenge posed by full-length barcodes; i.e., problems with reliably obtaining amplicons from museum specimens with degraded DNA (e.g. Hajibabaei et al. 2006). We thus submit that it is time to use empirical evidence to determine at which length and in which position mini-barcodes are as effective for pre-sorting specimens into putative species as full-length barcodes.

### Mini-barcodes

Barcodes that are shorter than the full-length barcode are often referred to as “mini-barcodes”. They are obtained with primers that amplify shorter subsets of the original barcode region and have three key advantages. Firstly, mini-barcodes are predicted to amplify more readily when the DNA in the sample is degraded. This was confirmed by several studies comparing amplification success rates for mini- and full-length barcodes for the same DNA template (Hajibabaei et al. 2006; Meusnier et al. 2008; Hajibabaei and McKenna 2012). This property makes mini-barcodes the preferred choice for barcoding museum specimens that had been stored under suboptimal conditions for decades. The ease of amplification is also one of the reasons why mini-barcodes are the default for metabarcoding projects that rely on environmental DNA with widely varying DNA quality [e.g., gut content, faeces (Deagle et al. 2006)].

A second benefit of mini-barcodes is that they can be sequenced at low cost on short-read sequencing platforms (e.g., Illumina). The maximum amplicon lengths that such platforms can accommodate is ∼450-bp including primer length (using 250-bp paired-end sequencing libraries), which necessitates the amplification and sequencing of two overlapping regions to recover the entire 658-bp barcode (Shokralla et al. 2015b). However, given the additional time and resources required, it begs the question if there is much to be gained by investing in full-length barcodes. Full-length barcodes are often automatically assumed to improve performance but more data does not automatically yield better results. There is often a saturation point beyond which additional data have little impact on results while increasing consumable and manpower cost. Occasionally, more data can even be detrimental. For example, fast-evolving genes or third positions in protein-encoding genes can worsen the results of phylogenetic analyses addressing the relationships between old clades. Note that this could also be a concern for barcodes, because sequence variability changes across the Folmer region of COI (Roe and Sperling 2007; Pentinsaari et al. 2016). Overall, it is important that one should use empirical data to determine how long barcodes have to be in order to be informative.

Unfortunately, the performance of mini-barcodes remains insufficiently tested despite their ubiquitous use in metabarcoding. The existing tests suffer from lack of scale (the largest study includes 6695 barcodes for 1587 species: Meusnier et al. 2008) or taxonomic scope (usually only 1-2 family-level taxa: e.g. Hajibabaei et al. 2006; Yu and You 2010). Furthermore, the tests yielded conflicting results. Hajibabaei et al. (2006) found high congruence with the full-length barcode when species are delimited based on mini-barcodes and Meusnier et al. (2008) find similar BLAST identification rates for mini-barcodes and full-length barcodes in their *in silico* tests. However, Yu and You (2010) conceded that mini-barcodes may have worse accuracy despite having close structural concordance with the full-length barcode. In addition, Sultana et al. (2018) concluded that the ability to identify species is compromised when the barcodes are too short (<150-bp), but it remained unclear at which length/position mini-barcodes stop performing poorly. Furthermore, published tests of mini-barcodes compare their performance to results obtained with full-length barcodes. All conflict is then considered evidence for the failure of mini-barcodes, even though the Folmer region varies in nucleotide variability (Roe and Sperling 2007) and conflict alone does not settle which result is correct. Lastly, the existing tests of mini-barcodes do not include a sufficiently large number of different mini-barcodes in order to be able to detect positional and length effects across the 658-bp barcode region.

Here, we address the lack of scale in previous tests by including 20 empirical studies covering 5500 species (ca. 30,000 barcodes) and Genbank data for 20,673 species (ca. 98,000 barcodes). We test a large number of different mini-barcodes by applying a sliding window approach to generate mini-barcodes of different lengths (100, 200, 300-bp window length, 60-bp intervals) and compare the results to the performance of nine mini-barcodes with established primers (mini-barcode length: 94 – 407-bp). The taxonomic scope of our study is broad and includes a wide variety of metazoans ranging from earthworms to butterflies and birds. Lastly, we do not assume that mOTUs based on full-length barcodes are more accurate than those obtained with mini-barcodes. Instead, we assess whether mOTUs obtained with different-length barcodes have different levels of congruence with morphospecies; i.e. morphology is treated as a constant and we only test whether barcode length and/or position influence the number of morphospecies that are recovered. All specimens that are mis-sorted based on morphology (e.g. due to the presence of cryptic species) will be equally mis-sorted for the full-length barcodes and the mini-barcodes that are derived from the former via shortening; i.e., misidentified specimens become noise. What we assess is whether there is an additional loss of congruence between barcodes and morphology as the barcode length shrinks or mini-barcodes are in a different position within the Folmer region of COI.

We also compare the performance of different species delimitation methods. There has been substantial interest in developing algorithms for mOTU estimation, leading to the emergence of various species delimitation algorithms over the past decade (e.g. objective clustering: Meier et al. 2006; BPP: Yang and Rannala 2010; jmOTU: Jones et al. 2011; ABGD: Puillandre et al. 2012; BINs: Ratnasingham and Hebert 2013; PTP: Zhang et al. 2013; etc.). For the purposes of this study, we selected three algorithms that represent distance- and tree-based methods: objective clustering, Automatic Barcode Gap Discovery (ABGD) and Poisson Tree Process (PTP). Objective clustering utilizes an *a priori* distance threshold to group sequences into clusters, ABGD groups sequences into clusters based on an initial prior and recursively uses incremental priors to find stable partitions, while PTP utilizes the branch lengths on the input phylogeny to delimit species units. Note that PTP is frequently applied to trees derived from barcode data (e.g. Ermakov et al. 2015; Han et al. 2016; Hollatz et al. 2016). There are numerous additional techniques for species delimitation, but most require multiple markers and thus do not scale easily to millions of specimens.

In addition to species delimitation, we also examine the utility of mini-barcodes for species identification, which is the original use of DNA barcodes (Hebert et al. 2003). Species identification with barcodes is particularly valuable for the detection of specific species and community characterization via metabarcoding studies (Ficetola et al. 2008; Lim et al. 2016; Morinière et al. 2019). Most of these studies tend to involve poor quality genetic material (i.e., gut content, faecal matter, environmental DNA) and hence it would be valuable to know whether short markers can be reliably used to assign species names to unidentified mini-barcodes. We here test the performance of the barcodes using “best close match”, which considers those sequences correctly identified when they match conspecific barcodes within a predefined threshold (Meier et al. 2006).

## Materials & Methods

### Dataset selection, alignment, excising in silico mini-barcodes

We surveyed the barcoding literature in order to identify 20 recent publications from 2007 – 2017 that cited the original barcode paper by Hebert et al. (2003) and met the following criteria: 1) have specimens where the barcoded specimens were pre-sorted/identified based on morphology and 2) the dataset had at least 500 specimens with *cox1* barcodes >656-bp (Table S1a). In addition, we compiled six additional datasets based on sequences from GenBank for species-rich metazoan taxa: Actinopterygii, Arachnida, Coleoptera, Diptera, Hymenoptera and Lepidoptera. The clade names were used for taxonomy searches in NCBI and all mitochondrial nucleotide sequences of length 657-1000 bp were downloaded. To these, we added annotated *cox1* sequences from mitochondrial genomes (mitochondrial sequences of length 10,000-20,000 bp). Subsequently, we filtered for the *cox1* gene using an alignment to a reference full-length *cox1* barcode with MAFFT’s --add function (Katoh and Standley 2013), where any non-target sequences are easily identified by large indels and discarded. We then removed all sequences that were shorter than 657-bp, introduced indels in the multiple sequence alignment, were not identified to species, or included ambiguous species names. Four of the datasets (Actinopterygii: 21,390 barcodes, Coleoptera: 44,403 barcodes, Diptera: 56,344 barcodes, Lepidoptera: 198,441 barcodes), were too large for a full evaluation using all species delimitation methods. We therefore randomly selected 20,000 sequences (=RAND() in Microsoft Excel) for further analysis. These six datasets comprised 97,758 barcodes for 20,673 morphospecies after subsampling (Table S1b).

The barcode sequences were downloaded from BOLDSystems or NCBI GenBank and aligned with MAFFT v7 (Katoh and Standley 2013) with a gap opening penalty of 5.0. Using a custom Python script, we generated three sets of mini-barcodes along a “sliding window” for the 20 empirical datasets. They were of 100-, 200- and 300-bp lengths. The first iteration begins with the first base pair of the 658-bp barcode and the shifting windows jump 60-bp at each iteration, generating ten 100-bp windows, eight 200-bp windows and six 300-bp windows. Additionally, we identified nine mini-barcodes with published primers within the *cox1* Folmer region (Fig. 1 & Table S2). These mini-barcodes have been repeatedly used in the literature for a broad range of taxa. The primers for the various mini-barcodes were aligned to the homologous regions of each dataset with MAFFT v7 --addfragments (Katoh and Standley 2013) in order to identify the precise position of the mini-barcodes within the full-length barcode. The mini-barcode subsets from each barcode were then identified after alignment to full-length barcodes to mark the start and end-point of the amplicon. Note that most of the published primers are in the 5’ region of the full-length barcode. The nine mini-barcodes with published primers were assessed for the 20 empirical and for the six GenBank datasets.

**Figure 1.**
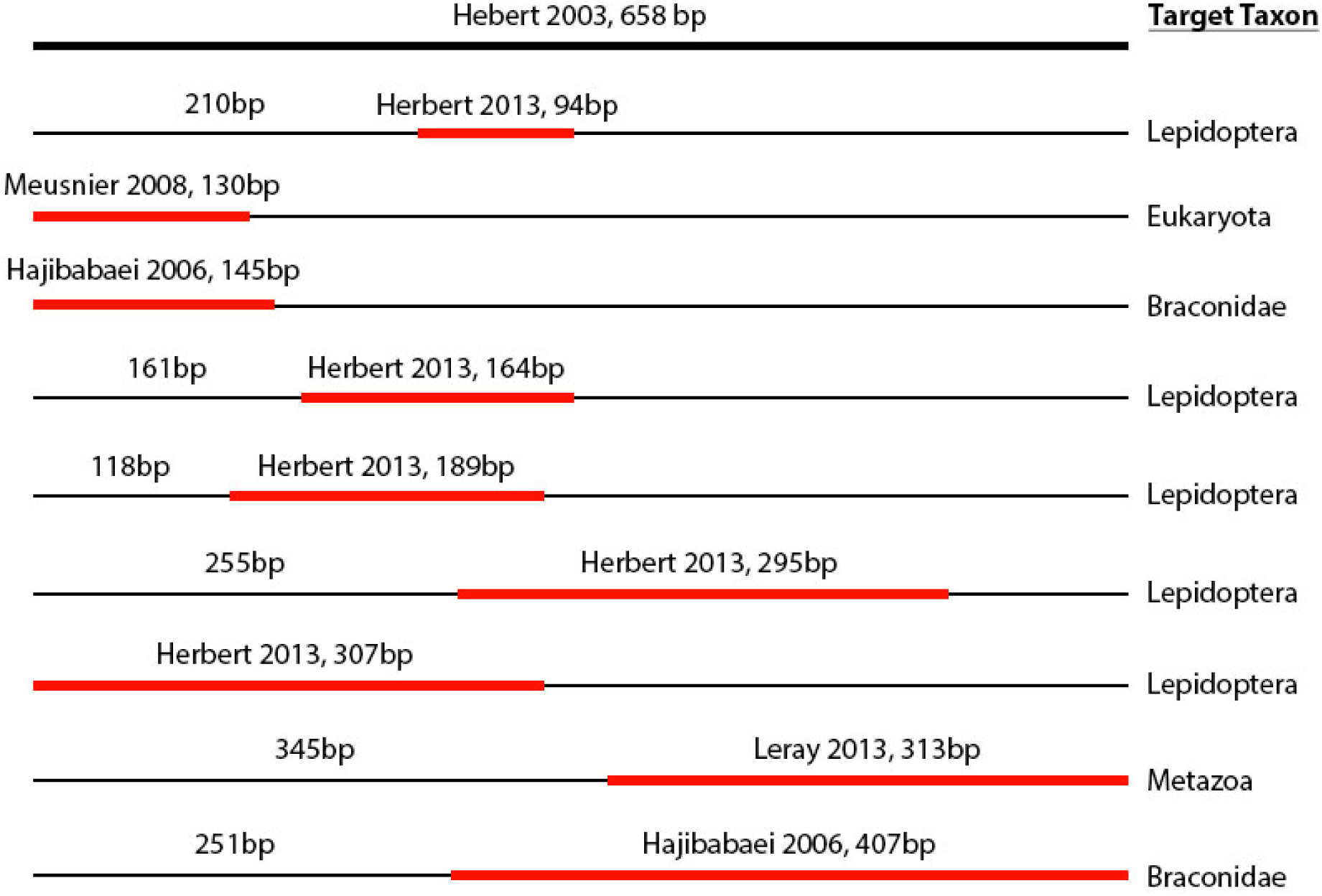
Positions of the mini-barcodes with established primers assessed in this study.

### Species delimitation

Mini-barcodes and full-length barcodes were clustered into putative species using three species delimitation algorithms: objective clustering (Meier et al. 2006), ABGD (Puillandre et al. 2012) and PTP (Zhang et al. 2013). For objective clustering, the mOTUs were clustered at 2 – 4% uncorrected p-distance thresholds (Srivathsan and Meier 2012) using a Python script which re-implements the objective clustering of Meier et al. (2006) and allows for batch processing. The p-distance thresholds selected are the typical distance thresholds used for species delimitation in the literature (Meier et al. 2006, 2016; Ratnasingham and Hebert 2013). The same datasets were also clustered with ABGD (Puillandre et al. 2012) using the default range of priors and with uncorrected p-distances, but the minimum slope parameter (-X) was reduced in a stepwise manner (1.5, 1.0, 0.5, 0.1) if the algorithm could not find a partition. We then considered the ABGD clusters at priors P=0.001, P=0.01 and P=0.04. The priors (P) refer to the maximum intraspecific divergence and function similarly to p-distance thresholds at the first iteration. In the ABGD algorithm, they are refined by recursive application. Lastly, in order to use PTP, we generated maximum likelihood (ML) trees in RAxML v.8 (Stamatakis 2014) via rapid bootstrapping (-f a) under the GTRCAT model. For the 20 empirical datasets, the best tree generated for each dataset was then used for species delimitation with PTP (Zhang et al. 2013) under default parameters. For the six much larger clade-based datasets, we used mPTP (Kapli et al. 2017) with the --single parameter as the original algorithm took too long to yield results. In order to reduce computational time for RAxML and PTP, haplotype representatives were used for the six Genbank dataset instead of the entire dataset. The remaining barcodes with identical haplotypes were then mapped back into the same mOTU clusters using their respective haplotypes.

### Performance assessment

The performance of mini-barcodes was assessed using morphospecies as an external arbiter. Species-level congruence was quantified using match ratios between molecular and morphological groups (Ahrens et al. 2016). The ratio is defined as 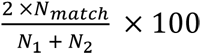, where N_match_ is the number of clusters identical across both mOTU delimitation methods/thresholds (N_1_ & N_2_). Consolidated match ratios are derived by first summing relevant all numerator and denominator values before calculating the ratio. Incongruence between morphospecies and mOTUs is usually caused by a few specimens that are assigned to the “incorrect” mOTUs. Conflict at the specimen-level can thus be quantified as the number of specimens that are in mOTUs that cause conflict with morphospecies.

In order to test whether barcode length is a significant predictor of congruence, MANOVA tests were carried out in R (R Core Team 2017) with “match ratio” (species-level congruence) as the response variable and “dataset” and “mini-barcode” as categorical explanatory variables. We found that most of the variance in our study was generated by the variable “dataset” (P < 0.05 in MANOVA tests). Given that we were interested in the effect of barcode length and position, “dataset” was subsequently treated as a random effect and “mini-barcode” as the explanatory variable (categorical) in a linear mixed effects model (R package *lme4*: Bates et al. 2014). The *emmeans* R package (Lenth 2018) was then used to perform pairwise post-hoc Tukey tests between mini- and full-length barcodes so as to assess whether either barcode was performing significantly better/worse. To compare the differences in performance between objective clustering, ABGD and PTP, ANOVA tests were performed in R. After which, pairwise Tukey tests were used to determine which species delimitation method was responsible for significant differences. Lastly, in order to explore the reasons for positional effects, the proportion of conserved sites for each mini-barcode was obtained using MEGA6 (Tamura et al. 2013).

Match ratios indicate congruence at the species level, but it is also important to determine how many specimens have been placed into congruent units. Note that species- and specimen-level congruence are only identical when all mOTUs are represented by the same number of specimens. However, specimen abundances are rarely equal across species and hence match ratio is insufficient at characterizing congruence between mOTUs and morphospecies. It is straightforward to determine the number of congruent specimens as follows:

1. Congruence Class I specimens: If ***A*** = ***B*** then number of congruent specimens is Nc_1_ = |*A*| OR |*B*|. Lack of congruence is caused by morphospecies that are split, lumped, or both split and lumped in the mOTUs. This means that a single mis-sorted specimen placed into a single, large-sized mOTU leads to all specimens in two mOTUs to be considered “incongruent” according to the criterion outlined above. Yet, most specimens are congruent and full congruence could be restored by re-assigning the mis-sorted specimen after re-examination. It is therefore also desirable to determine the number of specimens that require re-examination or, conversely, the number of specimens that would be congruent if one were to remove a few incongruent specimens. This number of specimens can be estimated by counting congruent specimens as follows:
2. Congruence Class II specimens: Specimens that are in split or lumped mOTUs relative to morphospecies. Here, the largest subset of congruently placed specimens can be determined as follows. If 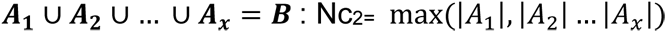
3. Congruence Class III specimens: This covers specimens in sets of clusters that are both split and lumped relative to morphospecies. Here, only those specimens are considered potentially congruent that (1) are in one mOTU and one morphospecies and (2) combined exceed the number of the other specimens in the set of clusters. In detail, if 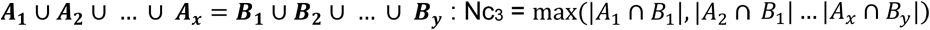 only if 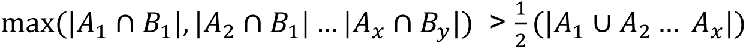. Note that these are approximate (over)estimates because we did not re-cluster specimens after the removal of the specimens that were causing incongruence in the original sets of clusters.

### Species identification

Using the full-length and mini-barcode sets corresponding to published primers, we assessed whether the barcodes were yielding conspecific matches according to “best close match” implemented in SpeciesIdentifier program (TaxonDNA: Meier et al. 2006). “Best close match” examines one barcode at a time. It considers it unidentified and then determines whether the best-matching reference sequence in the dataset is conspecific and within a user-defined distance threshold. The distance threshold used for all assessments was 2%, which corresponds to the best match ratio performance results for objective clustering in this study. Sequences with conspecific matches within 2% are considered correctly identified, while sequences that match allospecific sequences within 2% are considered incorrectly identified. Sequences without matches within 2% are considered unidentified and sequences that have both con- and allo-specific matches within the threshold are considered to have an ambiguous identification. Mixed effects models were again used to assess the performance of barcodes of different length. Similarly, pairwise Tukey tests were used to test for statistical significance with the proportion of correct matches as the response variable in order to determine whether the full-length barcode performed significantly better than the mini-barcodes.

### Amplification success rates for 313-bp and 658-bp amplicons

We tested amplification success rates for 47-48 specimens (∼1mm) from six Malaise trap samples of different age (2012 and 2018). The same genomic DNA was used for each specimen for the amplification of the full-length barcode and a mini-barcode of 313-bp length in two separate PCRs using different tagged primers: HCO2198 (5’-TAAACTTCAGGGTGACCAAAAAATCA-3’) and LCO1490 (5’- GGTCAACAAATCATAAAGATATTGG-3’) (658-bp; Folmer et al. 1994), mlCOIintF (5’-GGWACWGGWTGAACWGTWTAYCCYCC-3’) and jgHCO2198 (5’- TAIACYTCIGGRTGICCRAARAAYCA-3’) (313-bp; Leray et al. 2013). The PCR reagents used were 5.0µl Taq MasterMix (CW0682; CWBio, Beijing, China), 1.0µl RNase-free water, 0.5µl bovine serum albumin (1.0 mM), 0.5µl MgCl_2_, 1.0µl of each primer and 1.0µl of genomic DNA extract. The PCRs were run at 95°C for 5min, with 35 cycles of 94°C for 1min, 45°C for 2min and 72°C for 1min, and a final extension at 72°C for 5min. PCR amplification success was assessed using a 1% agarose gel, with the electrophoresis run at 90V for 30min. Band presence was tabulated and chi-squared performance testing was performed in R (R Core Team 2017).

## Results

### Evaluation of mini-barcode congruence with morphology

For species delimitation with objective clustering, we found that the 2% p-distance threshold yielded the highest congruence-levels across the datasets. It was hence used as the upper-bound estimator for assessing species- and specimen-level congruence (see supplementary materials for corresponding results for the 3 and 4% p-distances). For ABGD, average congruence was maximized for the prior P=0.001 and this prior was used in the main analysis (see supplementary material for results under P=0.01 and P=0.04). PTP does not require *a priori* parameter choices. Overall consolidated match ratio values range from 68 – 74% for these selected parameters (Fig. 3), with the 20 empirical datasets generally performing better than the six clade-based datasets (empirical/clade-based: OC at 2%: 81/72%; ABGD P=0.001: 76/69%; PTP: 76/66%). Overall match ratios for the other objective clustering and ABGD parameters tested range from 61 – 74% (Fig. S6).

The MANOVA tests performed on all treatments (species delimitation method and distance threshold/prior) indicated that the test variable “dataset” was responsible for much of the observed variance in our measure of congruence (“match ratio”). The choice of mini-barcode or algorithm for generating mOTUs was of secondary importance (Table S3). After accounting for “dataset”, we find that only mini-barcodes <200-bp perform significantly worse than full-length barcodes (Fig. 2). This is evident from the large number of significant differences (p < 0.05 & p < 0.001) in pairwise post-hoc Tukey tests applied to 100-bp mini- and 657-bp full-length barcodes. Only short barcodes (<100-bp) have a mean performance that is worse (<0 match ratio deviation) than the full-length barcode. Conversely, for mini-barcodes >200-bp, congruence with morphospecies does not differ significantly and is occasionally superior to what is observed for the full-length barcode. Across all datasets (20 empirical and 6 clade-based), species delimitation methods, and clustering parameters (Fig. 3 & S6), the full-length barcode have the highest congruence values in 49 of the 182 tests, but in 8 of these cases it is a tie and in an additional 19 some mini-barcodes are within 0.5%. Furthermore, there is no significant difference between the 200-bp and 300-bp mini-barcodes and the full-length barcode when objective clustering or PTP are used to estimate mOTUs. However, for all mOTU delimitation methods, the variance across datasets declines as the mini-barcode increases in length (Fig. 2). The results obtained for *in silico* mini-barcodes are broadly consistent with the performance of mini-barcodes corresponding to published primers: the mini-barcodes of 94-bp, 130-bp, and 145-bp lengths tend to perform worse than the longer mini-barcodes (Fig. 3). Congruence at the species level is similar to congruence at the specimen level (Tables 1 & S8) with some exceptions such as the performance improvements of short mini-barcodes, for example, for the “French Guiana earthworms” when grouped with objective clustering.

**Figure 2.**
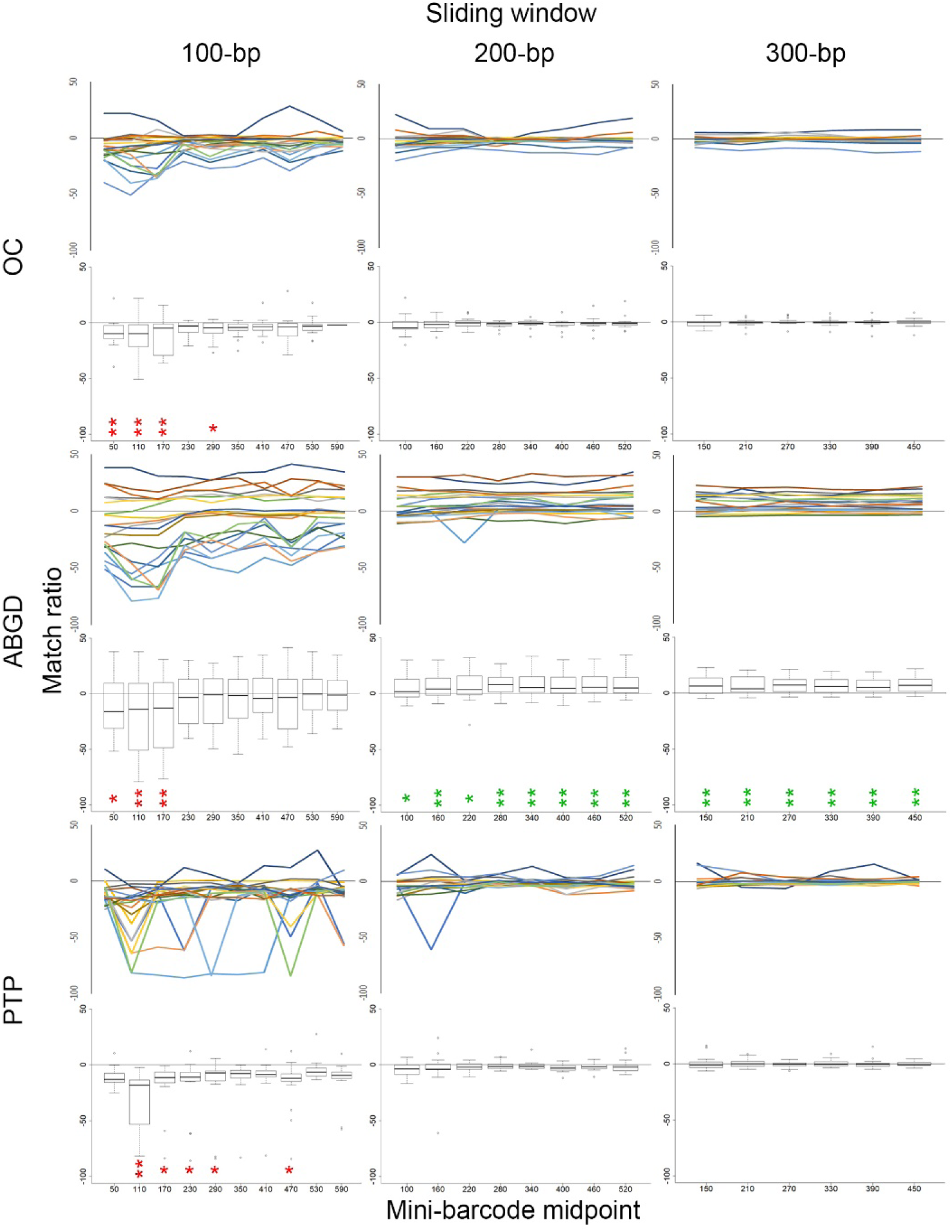
Performance of mini-barcodes along a sliding window (100-bp, 200-bp, 300-bp). Mini-barcode position is indicated on the x-axis and congruence with morphology on the y-axis. mOTUs were obtained with Objective Clustering (2%), ABGD P=0.001 prior), and PTP. Each line represents one data set while the boxplots summarise the values across datasets. Significant deviations from the results obtained with full-length barcodes are indicated with color-coded asterisks (* = p < 0.05; ** = p < 0.001; red = poorer and green = higher congruence with morphology).

The results obtained for the 20 empirical datasets are similar to what is found for the taxon-specific datasets consisting of sequences from GenBank. There are small differences in match ratios for barcodes >200-bp and only very short barcodes perform poorly. In many cases, the full-length barcode does not yield the best performance compared to the mini-barcodes. An exception is the Coleoptera where ABGD behaves erratically (Fig. 3). Short barcodes outperform long barcodes under the lowest prior (P=0.001), but this trend reverses for higher priors.

**Figure 3.**
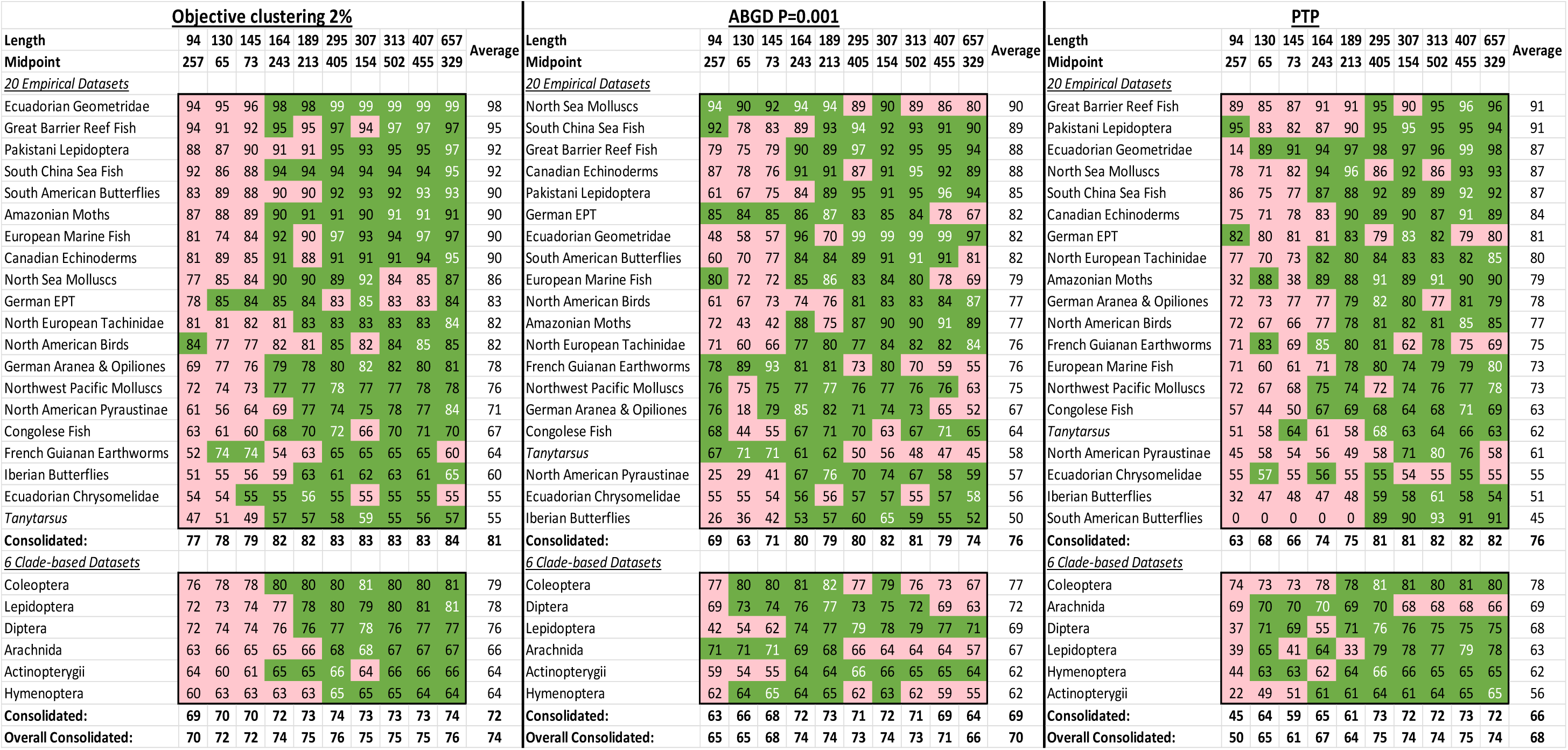
Match ratios across three different species delimitation methods. Mini-barcodes (columns) are sorted by primer length while the 20 empirical datasets (rows) are sorted by average match ratio. The six clade-specific datasets from GenBank are grouped below. Match ratios above average are coloured green, while those below average are in pink. The highest match ratio values are indicated with white text.

With regard to specimen-based congruence, we focused on the mini-barcodes with published primers that are >200-bp. For the 20 empirical datasets, approximately three quarters of all specimens are in the “Congruence Class I” (Tables 1 & S8); i.e., their placement is congruent between mOTUs and morphospecies (Average/Median: OC at 2%: 77/77%; ABGD P=0.001: 74/74%; PTP: 77/77%). The remaining specimens are placed in mOTUs that are split, lumped, or split and lumped. The number of specimens that are responsible for splitting and lumping are classified as Congruence Class II and III specimens (see Materials and Methods). Overall, fewer than 10% of the specimens fall into these categories (Table 1: see Class II specimens across species delimitation methods); i.e., a fairly small number of specimens have to be studied when addressing conflict between morphospecies and mOTUs. For the six clade-based datasets, there is generally lower specimen congruence (Class I Average/Median: OC at 2%: 57/57%; ABGD P=0.001: 53/54%; PTP: 57/57%), although this improves to ca. 75% at Class II and ca. 80% at Class III. Note that a single mis-identified *D. melanogaster* could result in all correctly identified specimens of this species to be considered “incongruent” because they are no longer member of a congruent mOTU.

**Table 1.** Proportion of specimens congruent between morphospecies and mOTU clusters (20 empirical datasets/6 clade-based datasets) under the three stringency classes. Values in brackets represent an upper bound estimate of number of specimens causing conflict.

| <u>Objective clustering, 2% p-distance</u> |  |  |  |  |  |  |  |
| --- | --- | --- | --- | --- | --- | --- | --- |
| Length | 295 | 307 | 313 | 407 | 657 | Average | Median |
| Midpoint | 405 | 154 | 502 | 455 | 329 |  |  |
| Class I | 77/57%<br>(7475/41367) | 76/56%<br>(7511/42052) | 77/57%<br>(7275/40857) | 77/57%<br>(7246/40849) | 77/58%<br>(7118/40327) | 77/57%<br>(7325/41090) | 77/57%<br>(7372/40941) |
| Class II | 90/75%<br>(3283/23846) | 90/75%<br>(3333/24794) | 90/75%<br>(3131/23554) | 90/76%<br>(3078/23304) | 90/76%<br>(3006/23349) | 90/75%<br>(3166/23769) | 90/76%<br>(3112/23469) |
| Class III | 91/82%<br>(3021/17875) | 91/81%<br>(2954/18981) | 91/82%<br>(2822/17666) | 91/82%<br>(2813/17511) | 91/82%<br>(2759/17509) | 91/82%<br>(2874/17908) | 91/82%<br>(2862/17745) |
| <u>ABGD, P=0.001</u> |  |  |  |  |  |  |  |
| Length | 295 | 307 | 313 | 407 | 657 | Average | Median |
| Midpoint | 405 | 154 | 502 | 455 | 329 |  |  |
| Class I | 75/55%<br>(7916/42576) | 76/55%<br>(7685/42403) | 75/55%<br>(8169/42787) | 73/53%<br>(8583/45131) | 69/47%<br>(9833/49848) | 74/53%<br>(8437/44549) | 74/54%<br>(8126/43386) |
| Class II | 89/75%<br>(3305/24555) | 89/75%<br>(3442/24541) | 89/75%<br>(3336/24267) | 89/74%<br>(3526/25153) | 88/72%<br>(3901/26691) | 89/74%<br>(3502/25041) | 89/74%<br>(3382/24625) |
| Class III | 91/81%<br>(2981/18303) | 91/81%<br>(2906/18509) | 90/81%<br>(2984/18352) | 90/80%<br>(3041/18978) | 89/78%<br>(3482/20860) | 90/80%<br>(3079/19000) | 91/81%<br>(2927/18494) |
| <u>PTP</u> |  |  |  |  |  |  |  |
| Length | 295 | 307 | 313 | 407 | 657 | Average | Median |
| Midpoint | 405 | 154 | 502 | 455 | 329 |  |  |
| Class I | 76/57%<br>(7643/40692) | 76/56%<br>(7744/42608) | 77/57%<br>(7369/41588) | 78/57%<br>(7130/41056) | 78/58%<br>(7207/40234) | 77/57%<br>(7419/41236) | 77/57%<br>(7318/40839) |
| Class II | 89/75%<br>(3322/23946) | 89/75%<br>(3549/24598) | 90/75%<br>(3230/23857) | 90/76%<br>(3109/23583) | 90/76%<br>(3127/22912) | 90/75%<br>(3267/23779) | 90/76%<br>(3198/23659) |
| Class III | 90/81%<br>(3091/18208) | 90/81%<br>(3334/19189) | 91/82%<br>(2926/18179) | 91/81%<br>(2813/18968) | 91/82%<br>(2859/17317) | 91/81%<br>(3005/18372) | 91/81%<br>(2937/18193) |

### Species delimitation method

When the performance of the three different clustering methods was compared, significant differences (p < 0.05 in ANOVA test) were found only for 100-bp mini-barcodes (Fig. 4). Here, pairwise post-hoc Tukey tests find that objective clustering performs significantly better than the other delimitation methods (p < 0.001) while ABGD and PTP do not differ significantly (p = 0.88) but behave erratically for short mini-barcodes (Fig. 2).

**Figure 4.**
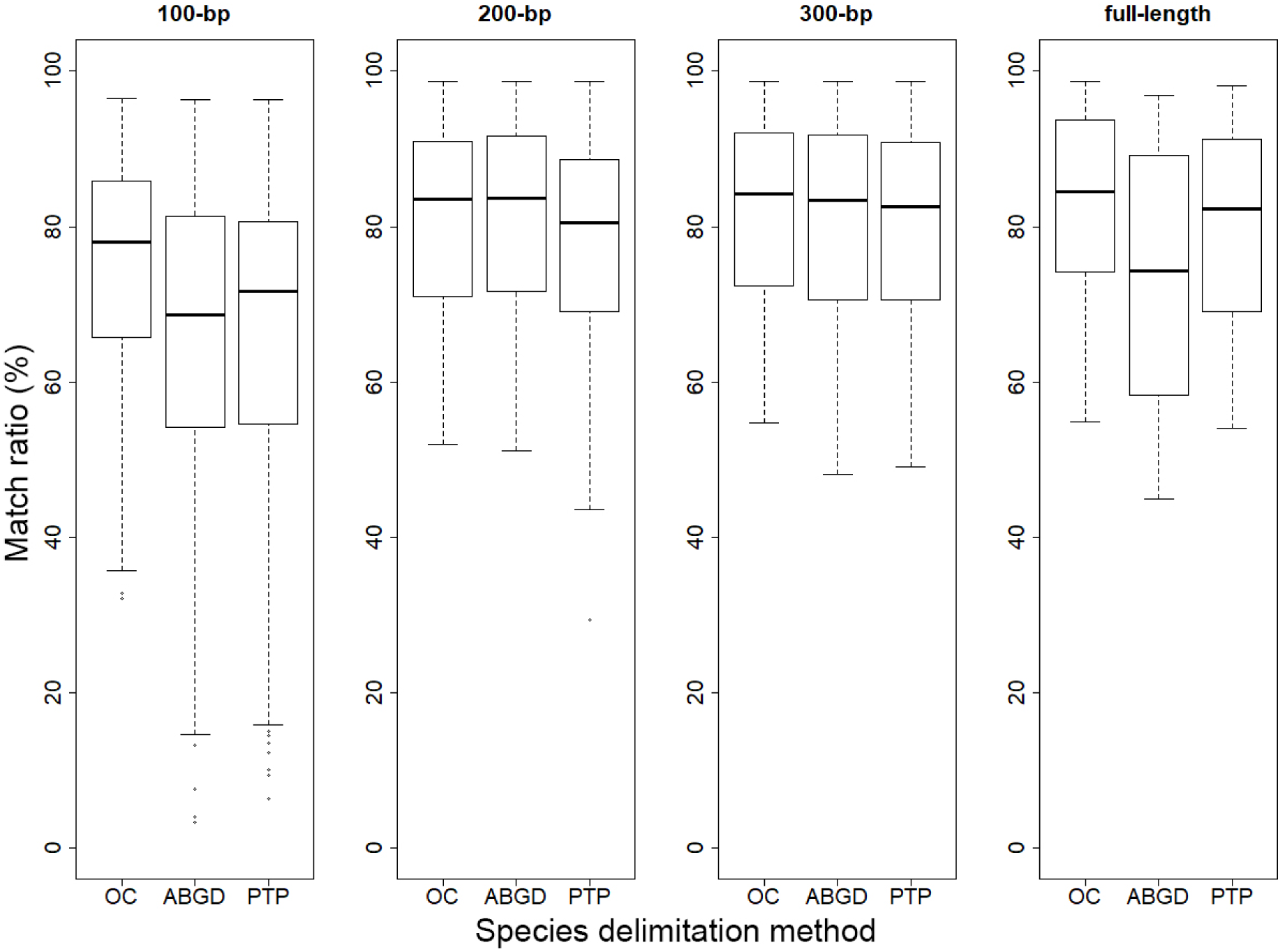
Comparison of species delimitation methods for full-length and mini-barcodes generated by “sliding windows” (100-bp, 200-bp, 300-bp).

### Positional effects

Mini-barcodes situated at the 5’ end of the full-length barcode perform somewhat worse than those situated at the middle or at the 3’ end (Fig. 2). For example, the 100-bp mini-barcodes at the 5’ end perform poorly for objective clustering (mini-barcode midpoints at 50, 110 & 170-bp), ABGD (mini-barcode midpoints at 110 & 170-bp) and PTP (mini-barcode midpoint at 110-bp). This effect is, however, only statistically significant when the mini-barcodes are very short (100-bp). This positional effect is observed across all species delimitation techniques. Note that the 5’ end of the full-length barcode appears to contain a large proportion of conserved sites, particularly around the 170-bp and 230-bp midpoint of the 100-bp mini-barcode (Fig. 5).

**Figure 5.**
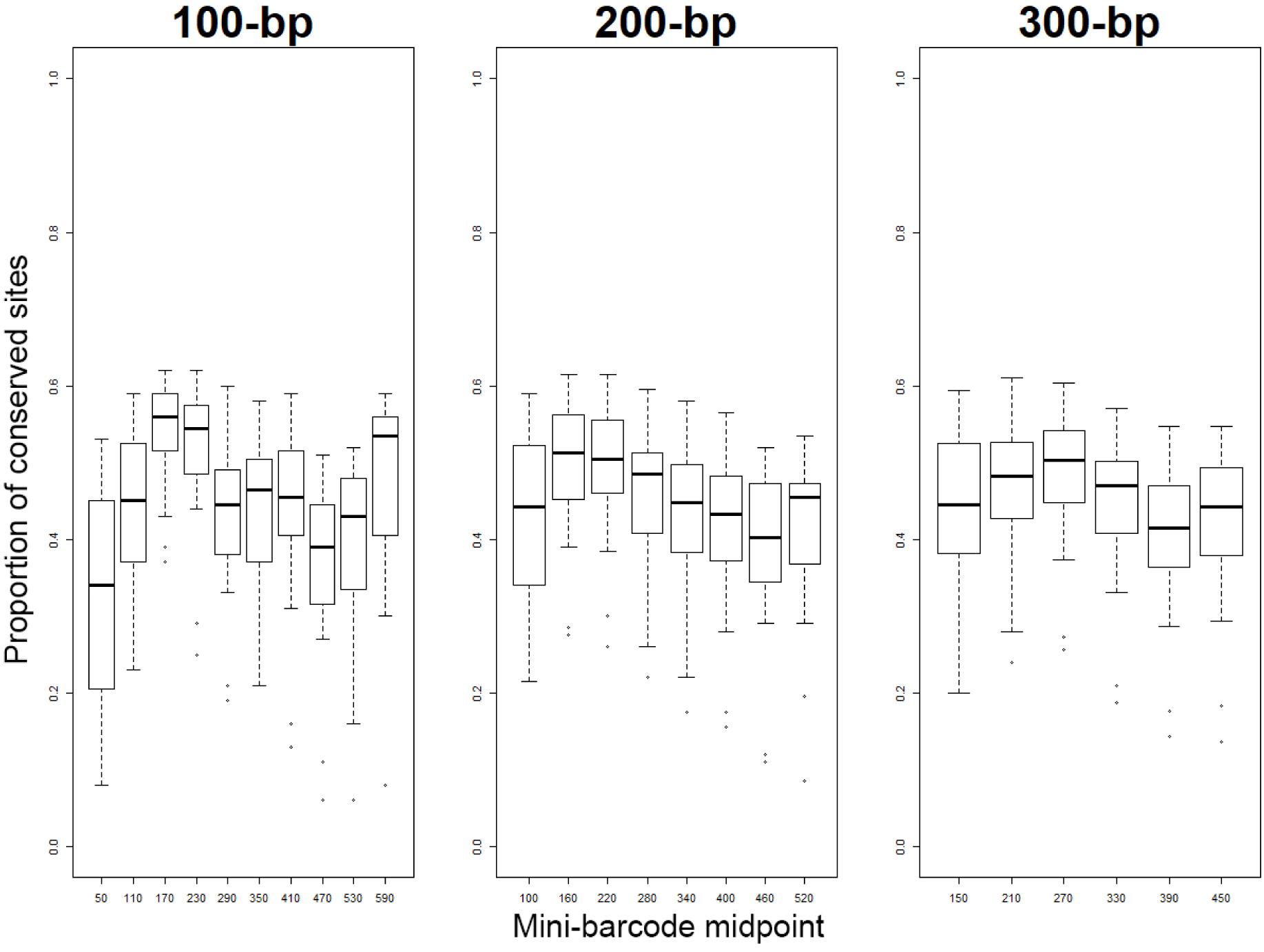
Proportion of conserved sites along the full-length barcode (sliding windows of 100-bp, 200-bp, 300-bp).

### Species identification performance

When comparing the suitability of full-length and mini-barcodes for identification purposes, we find that full-length barcodes frequently have the highest number of correct matches (Table 2), but the differences are only statistically significant for short mini-barcodes <200-bp: 94-bp: p = <0.0001, 130-bp: p = <0.0001, 145-bp: p = <0.0001, 164-bp: p = 0.0243. All the remaining mini-barcodes: (189 – 407-bp) have non-significant differences. Note that this is not due to scale because we are using fairly large datasets here (p = 0.0691 – 0.9996). However, for many datasets (e.g., Northwest Pacific Molluscs, North American Birds, etc.), the proportion of ambiguous matches is higher for very short mini-barcodes (94-bp – 164-bp).

**Table 2.** Proportion of correct/incorrect/ambiguous/unidentified specimens in the “best close match” analysis.

| Dataset | Published Primers |  |  |  |  |  |  |  |  |  |
| --- | --- | --- | --- | --- | --- | --- | --- | --- | --- | --- |
|  | 94 | 130 | 145 | 164 | 189 | 295 | 307 | 313 | 407 | 657 |
| Great Barrier Reef Fish | 85/1/0/15 | 83/1/3/13 | 83/1/2/14 | 85/0/0/14 | 86/1/0/13 | 86/0/0/14 | 85/1/0/14 | 86/0/0/14 | 86/0/0/14 | 86/0/0/14 |
| South China Sea Fish | 93/0/1/6 | 91/1/3/6 | 91/1/3/6 | 93/0/1/6 | 93/0/1/6 | 94/0/0/6 | 93/0/1/6 | 93/1/1/6 | 94/1/0/6 | 94/0/0/6 |
| North Sea Molluscs | 90/0/0/10 | 91/0/0/9 | 92/0/0/8 | 92/0/0/8 | 92/0/0/8 | 92/0/0/8 | 92/0/0/8 | 91/0/0/9 | 91/0/0/9 | 92/0/0/8 |
| Canadian Echinoderms | 95/0/0/5 | 95/0/0/4 | 95/0/0/5 | 96/0/0/4 | 96/0/0/4 | 95/0/0/5 | 96/0/0/4 | 95/0/0/5 | 96/0/0/4 | 96/0/0/4 |
| Ecuadorian Geometridae | 84/0/0/15 | 85/1/0/13 | 85/0/0/14 | 85/0/0/14 | 85/0/0/14 | 86/0/0/14 | 86/0/0/14 | 86/0/0/14 | 86/0/0/14 | 86/0/0/14 |
| German Aranea & Opiliones | 88/0/5/7 | 89/0/5/5 | 89/0/5/6 | 90/1/4/5 | 90/0/4/5 | 92/0/2/6 | 91/1/3/5 | 92/0/2/5 | 92/1/2/5 | 93/1/2/5 |
| European Marine Fish | 92/0/7/1 | 91/0/7/1 | 92/0/6/2 | 96/0/3/1 | 96/0/3/1 | 97/0/1/1 | 97/0/1/1 | 96/0/2/1 | 97/0/1/1 | 98/0/1/1 |
| South American Butterflies | 89/0/4/7 | 91/1/2/6 | 92/0/1/6 | 92/0/1/6 | 93/0/0/6 | 93/0/0/6 | 94/0/0/6 | 94/0/0/6 | 94/0/0/6 | 94/0/0/6 |
| German EPT | 92/0/1/7 | 93/0/1/5 | 94/0/1/6 | 94/0/0/6 | 94/0/0/6 | 93/0/0/6 | 94/0/0/5 | 94/0/0/6 | 94/0/0/6 | 94/0/0/6 |
| Northwest Pacific Molluscs | 69/2/15/14 | 75/2/10/13 | 74/2/10/13 | 73/2/12/12 | 73/2/12/12 | 77/2/8/12 | 77/2/9/12 | 76/3/9/12 | 77/3/8/12 | 78/4/6/12 |
| Amazonian Moths | 62/2/1/36 | 63/3/1/34 | 63/3/1/34 | 62/2/1/35 | 63/2/1/35 | 63/1/1/35 | 63/2/1/34 | 63/1/1/35 | 63/1/1/35 | 63/2/1/34 |
| North American Birds | 77/1/11/11 | 72/2/15/10 | 73/2/15/11 | 81/2/7/10 | 80/1/9/10 | 83/2/5/11 | 81/2/6/11 | 84/2/3/10 | 85/2/3/10 | 85/2/2/11 |
| French Guianan Earthworms | 96/0/0/4 | 97/0/0/3 | 97/0/0/3 | 96/0/0/4 | 96/0/0/4 | 97/0/0/3 | 97/0/0/3 | 97/0/0/3 | 97/0/0/3 | 96/0/0/4 |
| Pakistani Lepidoptera | 88/1/3/8 | 90/1/1/8 | 90/1/1/8 | 91/1/1/8 | 91/1/0/8 | 91/0/0/8 | 91/0/0/8 | 91/0/0/8 | 91/0/0/8 | 91/0/0/8 |
| *Tanytarsus* | 81/1/5/12 | 83/2/5/11 | 82/2/5/12 | 86/2/2/11 | 85/2/3/10 | 83/2/3/12 | 86/2/3/9 | 85/2/2/10 | 86/2/1/11 | 87/2/1/11 |
| North European Tachinidae | 64/2/14/19 | 65/3/16/17 | 65/3/15/17 | 66/3/13/18 | 65/3/15/17 | 67/3/11/18 | 68/3/12/17 | 68/3/10/18 | 69/3/10/18 | 71/4/8/17 |
| Congolese Fish | 80/1/10/9 | 78/1/11/9 | 79/1/11/10 | 84/2/6/7 | 86/2/5/7 | 85/2/5/8 | 86/2/4/8 | 86/3/4/7 | 86/3/4/7 | 88/3/2/7 |
| North American Pyraustinae | 93/1/4/3 | 94/1/4/1 | 96/1/2/1 | 96/1/1/2 | 97/0/0/2 | 97/0/0/3 | 98/1/0/2 | 97/0/0/3 | 97/0/0/2 | 98/2/0/2 |
| Ecuadorian Chrysomelidae | 74/0/1/25 | 76/0/1/23 | 76/0/1/23 | 76/0/1/23 | 76/0/1/24 | 75/0/1/24 | 76/0/1/23 | 75/0/1/24 | 75/0/1/24 | 75/0/1/24 |
| Iberian Butterflies | 77/0/22/1 | 85/0/15/0 | 86/0/13/0 | 87/0/12/0 | 89/0/10/0 | 93/0/6/0 | 94/1/5/0 | 94/0/5/0 | 96/0/4/0 | 97/1/3/0 |

### Amplification success rates

The 313-bp mini-barcode amplified significantly better than the full-length barcode (472/570: 82.8% vs. 361/570: 63.3%; chi-squared test: p = <0.001) (Fig. S9). Overall, the performance differences were greater for samples with overall low amplification success rates (but see sample 4822).

## Discussion

Accelerating species discovery and description are arguably foremost challenges for the systematics of the 21^st^ century because biodiversity is in steep decline while systematists are still grappling with establishing the size and distribution of the biodiversity that needs protection. Specimens for many undescribed species are already in the world’s natural history museums, but the samples need to be sorted to species-level before they become available for species identification/description and can be used for large-scale analyses of biodiversity patterns. Pre-sorting specimens with DNA barcodes is a potentially promising solution because it is scalable, can be applied to millions of specimens, and much of the specimen handling can be automated. However, in order for this approach to be viable, a sufficiently large proportion of the pre-sorted units need to accurately reflect species boundaries. Furthermore, the methods should be suitable for processing specimens that only yield degraded DNA. Mini-barcodes appear attractive on technical grounds because sequencing short amplicons is cheap (Wang et al. 2018: sequencing cost < 4 cents) and such amplicons have higher PCR amplification rates especially if the DNA template is degraded. The latter has been repeatedly discussed in the literature (Hajibabaei et al. 2006; Meusnier et al. 2008; Hajibabaei and McKenna 2012) and was also confirmed here by comparing identification success rates for a mini-barcode (313bp) and the full-full length barcode using the same template (Fig. S9).

However, the use of mini-barcodes for species-level pre-sorting would nevertheless have to be discouraged if full-length barcodes were to significantly outperform mini-barcodes with regard to the accuracy of sorting specimens into species. This is what we tested here and we find that mini-barcodes of moderate length (>200 bp) perform as well as full-length barcodes.

### The main source of variance of congruence: datasets

Overall, we here find that the average congruence between mOTUs and morphospecies is 80% for all barcodes >200-bp (median: 83%) in the 20 empirical datasets, with the median being higher (83%) because of a few outlier datasets with low congruence <65% (OC at 2%; ABGD P=0.001, PTP). Similar variance is also observed for the six clade-specific GenBank datasets. Both types of datasets were tested because GenBank data are less likely to be affected by the idiosyncrasies of individual study designs, but is known to contain misidentified sequences (Mioduchowska et al. 2018). We overall find that the results are nevertheless similar (Fig. 3 & S6) although the average congruence between mOTUs and morphospecies is somewhat lower for GenBank datasets (mean=72%, median=71% for all barcodes >200-bp across all three species delimitation methods and parameters). Full-length barcodes only have the highest congruence level in 13 of the 42 treatments, but in 10 of those, the second highest congruence level for a mini-barcode was only <0.5% lower. Mini-barcodes <200-bp perform poorly with the possible exception of those for Coleoptera where mini-barcodes <200-bp perform well under ABGD under two priors (Figs. 3 & S6: P=0.001, P=0.01).

Most studies assessing congruence levels between barcodes and morphology focus on species-level congruence. However, specimen-level congruence is equally important because the basic units in a museum collection or ecological surveys are specimens. The correct placement of specimens into species is thus important for systematists and biodiversity researchers alike given that the former would like to see most of the specimens in a collection correctly placed and the latter often need abundance and biomass information with species-level resolution. We find that for the 20 empirical datasets, 74-77% (median) of the ca. 30,000 specimens are assigned to species that are supported by molecular and morphological data even under the strictest conditions (Table 1). Overall, this is a very high proportion compared to species-level congruence obtained with species-level sorting by parataxonomists (Krell 2004).

The remaining ca. 25% of specimens are placed in mOTUs whose boundaries do not agree with morphospecies. One may initially consider this an unacceptably high proportion, but it is important to keep in mind that the misplacement of one specimen (e.g., due to a misidentification or contamination of a PCR product) will render two mOTUs incongruent; i.e., all specimens in these mOTUs will be considered incongruent and contribute to the 25% of “incongruently” placed specimens. Arguably, one may instead want to investigate how many specimens are causing the conflict. These are the specimens that should be targeted for re-investigation in reconciliation studies using additional data. The proportion across the 20 datasets in our study is fairly low and ranges from 9-11% (median) depending on which mOTU delimitation technique is used.

The six GenBank datasets however, have notably lower congruence-levels for specimens (Class I median: 54-57%), with the Actinopterygii (47-48% median) and Hymenoptera (45-47% median) having the poorest performance (Table S8). The overall lower congruence could be caused by the misidentification of a few specimens belonging to large mOTUs. This is supported by the greater difference between Class I and Class III for the clade-based datasets as compared to the empirical ones. Identifications from the empirical datasets are also more likely to be standardized within the study and consequently have fewer potential conflicts caused by differing taxon concepts associated with the same name (Franz 2005; Meier 2017).

Conflict between mOTUs and morphospecies can be caused by technical error or biology. A typical technical factor would be accidental misplacement of specimens due to lab contamination or error during morphospecies sorting. Indeed, the literature is replete with cases where mOTUs that were initially in conflict with morphospecies became congruent once the study of additional morphological characters let to the revision of morphospecies boundaries (e.g. Smith et al. 2008; Tan et al. 2010; Baldwin et al. 2011; Ang et al. 2017). But there are also numerous biological reasons why one should not expect perfect congruence between mOTUs and species. Lineage sorting, fast speciation, large amounts of intraspecific variability, and introgression are known to negatively affect the accuracy of DNA barcodes (Will and Rubinoff 2004; Rubinoff et al. 2006; Meier 2008). It is thus somewhat heartening that regardless of these issues, the final levels of congruence between morphospecies and DNA sequences are often quite high in animals (Ball et al. 2005; Cywinska et al. 2006; Renaud et al. 2012; Landi et al. 2014; Wang et al. 2018). This implies that pre-sorting specimens to species-level units based on barcodes is worth pursuing for many metazoan clades. This realization led to the proposal of the “reverse workflow” in Wang et al. (2018) who tested pre-sorting based on sequences for 4,000 ant specimens. They found that 86 of the 89 molecular operational taxonomic units (mOTUs) were congruent with morphospecies after reconciliation.

High levels of congruence levels are, however, not a universal observation across all of life. There are groups with widespread barcode sharing between species. Within Metazoa this is known for taxa such as Anthozoa (Huang et al. 2008) and is likely to be the default outside of Metazoa (e.g. Chase and Fay 2009; Hollingsworth et al. 2011). Indeed, congruence levels between mOTUs and morphospecies also vary widely in the 20 empirical and 6 Genbank datasets studied here (Fig. 3). Eighteen of the 26 datasets have congruence levels >75% and are arguably doing well. With regard to the remaining eight, there are four where the low levels may be due to the fact that they cover taxa or faunas that are comparatively poorly known and sorting with morphology may be prone to error (French Guiana Earthworms, Ecuadorian Chrysomelidae, chironomid midges in the genus *Tanytarsus*). For a 5^th^ data set, “Congolese fishes”, the authors also mention identification problems due to the lack of identification tools (Decru et al. 2016). However, these problems should be lacking for Iberian butterflies where the authors consider it likely that introgression and cryptic species are responsible for the lack of congruence (Dincă et al. (2015). Note, however, that these problems do not appear to affect the remaining Lepidoptera data set for which we observe high levels of congruence (5 data sets). The picture is similarly confusing for fish (Actinopterygii) where low congruence is observed for the Genbank dataset while marine fishes are doing well (3 datasets). Similarly, inconsistent patterns are observed for Hymenoptera and Arachnida.

We suspect that these inconsistent results are largely due to the lack of rigorous studies that establish congruence with confidence. Such studies should be based on datasets with dense taxon and geographic sampling where both morphological and DNA sequence information is obtained for all specimens. Subsequently, all specimens that cause conflict should be re-examined in order to determine the proportion of specimens that were initially misplaced based on morphology and the proportion that were initially misplaced based on DNA barcodes (=pre-sorting error). Lastly, one should establish the number of specimens that cannot be resolved because there is genuine conflict between the data sources. Such studies are lacking. This also affects the quality of the morphospecies sorting in the 26 datasets that are analzyed in our study. However, it should not affect our overall conclusions with regard to the effect of barcode length and position on congruence levels because morphospecies sorting errors affect both full-length barcodes and the mini-barcodes that were excised from the former.

But what is gained if all specimens are sorted based on morphology and barcodes? Using both sources of data would allow for higher quality work based on the principles of integrative taxonomy (Dayrat 2005; Schlick-Steiner et al. 2010), but it would not accelerate species discovery. Acceleration can only be achieved if either a subset of all specimens have to be studied with morphology or the morphological work requires less time. Both is the case when pre-sorting with barcodes is adopted. This is shown in Wang et al. (2018) and Srivathsan et al. (2019), who pre-sorted 4000 ant and 7000 phorid specimens, respectively, with barcodes. Checking for congruence between mOTUs and morpho-species was fast because the specimens had already been pre-grouped into putative species based on barcodes. In addition, determining the species limits between putatively closely related species was simplified because the taxon-specialist could identify closely related species/populations based on barcode distances. Lastly, for abundant species, only a subset of the specimens had to be studied with morphology. This subset could be selected based on genetic, temporal, and geographic gradients (Wang et al. 2018; Srivathsan et al. 2019); i.e., the highly-skilled specialists could target a small proportion of specimens for dissection and slide-mounting.

### Barcode length and species delimitation methods

We here tested the widespread assumption that mOTUs based on full-length barcodes are more reliable than those based on mini-barcodes (Burns et al. 2007; Min and Hickey 2007). If this assumption was confirmed, then the use of mini-barcodes for pre-sorting would have to be discouraged despite higher amplification success rates and lower cost. However, we find that the performance of *cox1* mini-barcodes with a length >200-bp do not differ substantially from the performance of full-length barcodes. In 19/140 of the empirical and 3/42 of the clade-based dataset treatments, the full-length barcode outperform all mini-barcodes, after discounting ties (8/140 and 0/42) and cases where the difference between the next highest congruence is <0.5% (9/140 and 10/42). Indeed, compared to the dataset effect, the choice of barcode length is largely secondary. Note that this conclusion is robust across 26 datasets and holds for different clustering algorithms.

We also find that the choice of species delimitation algorithm matters little for mini-barcodes longer than 200-bp (Fig. 4). This is fortunate as objective clustering and ABGD algorithms are less computationally demanding than PTP, which necessitates the reconstruction of ML trees. However, there are some exceptions. Firstly, when mini-barcodes are extremely short (∼100-bp), ABGD and PTP tend to underperform relative to objective clustering. PTP’s poor performance for 100-bp mini-barcodes is not surprising given that it relies on tree topologies which cannot be estimated with confidence based on so little data. ABGD’s poor performance is mostly observed for certain priors (e.g., P=0.04: Fig. S5 & S6). Under these priors, ABGD tends to lump most of the 100-bp barcodes into one or a few large clusters. Prior-choice also affects ABGD’s performance for full-length barcodes. Overall, ABGD does not perform well with very high priors (P = 0.04: Fig. S6 vs. P = 0.001: Fig. 2 & 3) and we conclude that the selection of the best priors and/or clustering thresholds remains a significant challenge for the study of largely unknown faunas that lack morphological or other gene information that can be used as an *a posteriori* criterion for selecting priors/thresholds. In the absence of these data, we would recommend the use of multiple methods and thresholds in order to distinguish robust from labile mOTUs that are heavily dependent on threshold- or prior-choices. The latter mOTUs should be targeted in reconciliation studies utilizing another type of data. As illustrated earlier, these reconciliation studies can focus on only 10-15% of all specimens that cause species-level conflict.

### Barcode length and species identification

Similar to our tests of species delimitation, we find that mini-barcodes >200-bp have no significant performance difference when compared with full-length barcodes. Full-length barcodes have only marginally – albeit statistically non-significantly – higher identification success rates than mini-barcodes >200-bp (Table 2: 0.00 – 4.15% difference, 0.37% median). This is consistent with the findings of Meusnier et al. (2008) who reported at least 95% identification success for 250-bp barcodes while full-length barcodes performed the best (97% success). Overall, our data suggest that mini-barcodes >200-bp are suitable for species identification. The poorer performance of shorter mini-barcodes (<200-bp) is largely due to an increase of ambiguous matches rather than incorrect matches. These mini-barcodes are less informative which is probably due to their short length and the placement of many short barcodes in a more conserved part of the Folmer region (see below).

### Positional effects

Overall, mini-barcodes at the 3’ end of the Folmer region outperform mini-barcodes at the 5’ end. This is consistent across all three species delimitation methods and was also noticed by Shokralla et al. (2015a) who concluded that mini-barcodes at the 5’ end have worse species resolution for fish species. This positional effect is apparent when match ratios are compared across a “sliding window” (Fig. 2). The lowest congruence with morphology is observed for 100-bp mini-barcodes with midpoints at the 50-bp, 110-bp and 170-bp marks. However, this positional effect is only significant when the barcode lengths are very short (<200-bp). Once the mini-barcodes are sufficiently long (>200-bp), there is no appreciable difference in performance, which is not surprising because sampling more nucleotides helps with buffering against regional changes in nucleotide variability.

These changes in nucleotide variability may have functional reasons related to the conformation of the Cox1 protein in the mitochondrion membrane. The Folmer region of Cox1 contains six transmembrane α-helices that are connected by five loops (Tsukihara et al. 1996; Pentinsaari et al. 2016). Pentinsaari et al. (2016) compared 292 Cox1 sequences across 26 animal phyla and found high amino acid variability in helix I and the loop connecting helix I and helix II (corresponding to position 1-102 of *cox1*), as well as end of helix IV and loop connecting helix IV and V (corresponding to positions ∼448-498). These regions of high variability are distant from the active sites and thus less likely to affect Cox1 function (Pentinsaari et al. 2016). This may lead to lower selection pressure and thus higher variability in these areas which could impact the performance of mini-barcodes for species delimitation.

### Accelerating biodiversity discovery and description

In our study we test whether species discovery with mini-barcodes is as accurate as discovery with full-length barcodes. However, the taxonomic impediment is also caused by the lack of taxonomic capacity for identifying specimens to species and describing new species. Fortunately, the species discovery step can also help with species identification because some specimens placed in mOTUs can be identified via barcode databases. Common species are here more likely to benefit because they are covered by the databases. This is illustrated by our recent work on dragon- and damselflies (Odonata), ants (Formicidae), and non-biting midges (Chironomidae) of Singapore (Baloğlu et al. 2018; Wang et al. 2018; Yeo et al. 2018). These studies document why it is important to distinguish between species- and specimen-level identification because the results differ widely. For odonates, BLAST-searches identified more than half of the 95 mOTUs and >75% of the specimens to species. The corresponding numbers for ants and midges were ca. 20% and 10% at mOTU-level, and 9% and 40% at the specimen-level; i.e., despite low mOTU identification success for midges, a fairly large number of specimens could be identified because very common species were already represented in a barcode database.

In addition, mOTUs are also useful for understanding biodiversity patterns because they can be readily compared across studies and borders (Ratnasingham and Hebert 2013). In contrast, a new species identified based on morphology usually remains unavailable to the scientific community until the descriptions are published unless associated molecular information is also present. This is a very significant difference because a large amount of downstream biodiversity analysis can be carried out based on newly discovered species regardless of whether they are were discovered with morphological or molecular means. However, cross-study comparisons are only straightforward for mOTUs because sequences are readily shared and compared. This allows for the study of species richness and abundances before taxa are described. Even studies covering different time periods can be carried out, which is increasingly important in the 21^st^ century.

### Conclusions

We here illustrate that mini-barcodes are as reliable for pre-sorting specimens into putative species as full-length barcodes. Mini-barcodes are also suitable for identifying unidentified barcodes to species based on barcode reference databases. We would thus argue that mini-barcodes should be preferred over full-length barcodes because (1) they can be obtained more readily for specimens that only contain degraded DNA (Hajibabaei et al. 2006) and (2) are much more cost-effective. In particular, we recommend the use of mini-barcodes >200-bp at the 3’ end of the Folmer region. It is encouraging that such mini-barcodes perform well across a large range of metazoan taxa. These conclusions are based on three species delimitation algorithms (objective clustering, ABGD and PTP) which, overall, have no appreciable effect on the performance of barcodes. If the DNA of the specimens is so degraded that very short mini-barcodes have to be obtained, we advise against the use of PTP and ABGD (especially with high priors) in order to reduce the likelihood that morphospecies are lumped.

## Acknowledgements

We would like to acknowledge Athira Adom and Jonathan Ho for data processing, Lee Wan Ting for performing the molecular work and Emily Hartop and Wendy Wang for commenting on the manuscript.

